# The SARS CoV-1 3a protein disrupts Golgi complex morphology and cargo trafficking

**DOI:** 10.1101/2021.04.19.440492

**Authors:** Rex R. Gonzales, Carolyn E. Machamer

## Abstract

Coronaviruses assemble by budding into the endoplasmic reticulum-Golgi intermediate compartment, but the pathway of egress from infected cells is not well understood. Efficient egress of infectious bronchitis virus (a gamma coronavirus, CoV) requires neutralization of Golgi pH by the envelope (E) protein. This results in reduced rates of cargo traffic and disrupts Golgi morphology, but it protects the spike protein from aberrant proteolysis. The severe acute respiratory syndrome (SARS) CoV-1 E protein does not disrupt the Golgi, however. We show here that in transfected cells, the ORF3a protein of SARS CoV-1 disrupts Golgi morphology, cargo trafficking and luminal pH. Unlike the infectious bronchitis virus E protein, these functions of the SARS CoV-1 3a protein appear to require its viroporin activity. Thus, neutralization of acidic compartments may be a universal feature of CoV infection, although different viral proteins and mechanisms may be used to achieve this outcome.

## Introduction

Coronaviruses are enveloped RNA viruses that assemble by budding into the endoplasmic reticulum-Golgi intermediate compartment. The envelope proteins (spike, S; membrane, M and envelope, E) are targeted to this compartment and virions bud into the lumen once the nucleocapsid-wrapped genome interacts with them [1]. The pathway for egress is not well understood, but given the size of the virions, the exocytic compartments must be modified to accommodate the large cargo [2]. Several membrane proteins encoded by CoVs have ion channel activity (called viroporins) that might be required for efficient virus egress [3, 4]. The CoV E protein is such a viroporin [5–7]. We previously studied a recombinant infectious bronchitis virus (IBV) with an E protein containing a mutated transmembrane domain. Release of infectious virus is reduced for this mutant and particles that are released have aberrantly proteolyzed S proteins, explaining the reduction in infectivity [8, 9]. In addition, overexpression of the wild-type IBV E protein, but not the mutant E protein, resulted in reduced cargo trafficking and fragmentation of the Golgi complex [10]. We recently demonstrated that the wild-type IBV E protein neutralized the Golgi complex in infected and transfected cells, whereas the mutant E protein did not [11]. Surprisingly, the ion channel activity of IBV E was not required for Golgi neutralization and disruption, since these phenotypes correlated with a mutant form of E that is predominantly monomeric [12]. Neutralization of the Golgi was required to protect the S protein from aberrant proteolysis and is likely the cause of Golgi disruption. The reduced cargo trafficking we observed is apparently an acceptable compromise to promote release of infectious virus.

IBV is a gamma CoV, and human CoVs fall into the alpha and beta genera. The human CoVs that cause severe disease, including SARS CoV-1 and -2, and the Middle East respiratory syndrome (MERS) virus, are all beta-CoVs. Whether secretory pathway modifications are universal requirements for CoV egress is not known. A recent paper on mouse hepatitis virus (MHV), a beta CoV, showed that endo-lysosomal compartments were neutralized in infected cells [13]. The authors proposed a novel egress pathway through lysosomes. The viral protein required for neutralization of endo-lysosomes in MHV-infected cells is not yet known.

Interestingly, we found that the SARS CoV-1 E protein did not possess Golgi disruption activity when overexpressed in cells [9]. We therefore investigated one of the accessory proteins, ORF 3a, since an earlier publication reported alteration of Golgi morphology when the protein was transiently expressed [14]. The 3a protein is quite different in structure than the small, single membrane spanning E protein, with three membrane spans. However, it does have cation channel activity and has been implicated in virus egress [15]. The Enjuanes group has shown that SARS CoV-1 lacking either E or 3a was viable, but virus lacking both could not be recovered [16]. We report here that SARS CoV-1 3a has similar Golgi disrupting activity to the IBV E protein, and the 3a protein from SARS CoV-2 appears to have a similar function. However, the mechanism by which SARS CoV-1 3a induces disruption of Golgi function differs from that of IBV E because viroporin activity is required.

## Materials and Methods

### Cells

Vero (African green monkey kidney) cells (ATCC CCL81) were maintained in Dulbecco’s modified minimal medium (Life Technologies) with 10% fetal bovine serum (Atlanta Biologicals) and normocin antibiotic (InVivoGen) at 37°C with 5% CO_2_. They were routinely screened for mycoplasma contamination.

### Plasmids

The 3a ORF from SARS CoV-1 (Tor2 strain) was originally obtained from the Institute for Genomic Research and subcloned into pBluescript (Stratagene) prior to transferring to pCAGGS-MCS [17] after PCR amplification with KpnI and XhoI restriction sites. SARS CoV-2 ORF 3a was subcloned from a plasmid kindly donated by Dr. Stephen Gould, Johns Hopkins Univ.) using KpnI and XhoI restriction sites. The pCAGGS plasmids encoding IBV E and SARS CoV-1 E have been previously described [10,18]. The SARS CoV-1 3aY109A mutant was generated by QuikChange mutagenesis (Stratagene) using the forward primer 5’-GCGCAATTTTTGTACCTGGCGGCCTTGATATATTTTC-3’ and the complementary reverse primer. Plasmids pCAGGS/SARS S and pCAGGS/SARS M were previously described [20], as was the pCAGGS/VSVG plasmid [21]. The pGnT1-pHorin and TGN-pHlorin plasmids [22] were kindly provided by Dr. Yasuke Maeda (Osaka University, Japan).

### Transfection

Vero cells were plated at 2×10^5^ per 35 mm dish on coverslips for indirect immunofluorescence microscopy, or 4×10^5^ in 6 well dishes for trafficking experiments. For the Golgi luminal pH measurement, Vero cells were plated at 6×10^5^ in 6 cm dishes. The following day, cells were transfected as follows. For immunofluorescence, 0.5 ug of SARS 3a or IBV E (with or without 0.2 ug of one of the pHlorin plasmids) by adding to 100 ul OPTI-MEM (Life Technologies) with diluted XtremeGene-9 (Roche), per the manufacturer’s instructions. For the trafficking experiments, 0.5 ug of 3a or E was mixed with 0.5 ug SARS CoV S or VSV-G. For flow cytometry, 6 cm dishes were transfected with 1.5 ug 3a or E plasmid with 0.5 ug GnT1-pHlorin, using 300 ul OPTI-MEM and 6 ul XtremeGene-9. Additional cells were transfected with the pHlorin plasmid alone to produce the standard curve for pH measurement.

### Antibodies

Rabbit polyclonal antibodies to IBV E, SARS CoV E, SARS CoV S and VSV-G were previously described [18–20, 23]. The SARS CoV-1 3a antibody was generated to a C-terminal peptide (CIYDEPTTTTSVPL-COOH) conjugated to KLH via the N-terminal cysteine residue and produced in rabbits by Covance Research Products. We determined specificity by immunoblotting and immunofluorescence using control and cells expressing SARS CoV-1 3a. SARS CoV-2 3a is identical in this epitope. Commercial primary antibodies were mouse anti-GM130 and mouse anti-p230 (BD Transduction Labs), and rabbit anti-giantin (Covance Research Products). Fluorescent secondary antibodies were from Life Technologies: Alexa Fluor 488 donkey anti-rabbit IgG, Alexa Fluor 568 donkey anti-mouse IgG, and Cy5 donkey anti-rabbit IgG.

### Indirect immunofluorescence microscopy and quantification of Golgi dispersion

Transfected cells were fixed at 18 h post-transfection. The protocol was exactly as described [11]. Images were acquired on an Axioskop microscope (Zeiss) equipped for epifluorescence using an Orca-03G CCD camera (Hamamatsu) and iVision software (BioVision Technologies). To quantify Golgi fragmentation, the area occupied by different Golgi resident proteins was determined. Images (acquired with the same parameters) were analyzed by outlining the stained Golgi elements after background subtraction using the trapezoid tool in ImageJ (Fiji), and the area occupied by each marker was quantified using the measure tool in the analysis menu. The total and integrated pixel intensity was also measured for each marker. The negative control was cells expressing IBV M, which is also Golgi targeted and the positive control was cells expressing IBV E. Analysis was performed in Prism GraphPad 8.0, with one-way ANOVA and post-hoc Tukey test. For all Golgi markers, including the peripheral membrane protein GM130, the total pixel intensities of stained elements in SARS CoV-1 3a-expressing cells were similar to those of cells expressing IBV M, suggesting the proteins were dispersed and not degraded.

### Cargo trafficking

Transfected cells were surface biotinylated to determine the level of cargo protein that had reached the plasma membrane. The lysates were also treated with endoglycosidase H (endo H) to determine the fraction that had moved past the medial Golgi by measuring oligosaccharide processing. For surface biotinylation, cells at 20 h post-transfection were rinsed in ice-cold PBS and then incubated at 0°C for 30 min with Hank’s buffered salt solution containing 0.5 ug/ml of EZ-link NHS-sulfo biotin (Pierce), with rocking. The biotin was quenched by incubation in PBS with 10 mM glycine for 10 min at 0°C, and cells were scraped, spun and lysed in 80 ul of 1% NP40, 150 mM NaCl, 10 mM Hepes pH 7.2 and 1x protease inhibitor cocktail (Sigma) on ice for 10 min, and clarified by centrifugation at 14K RPM for 10 min at 4°C. Ten percent of the lysate was reserved for endoH digestion. This portion of the lysate was denatured in 1x denaturation buffer (New England Biolabs) with incubation at 85°C for 15 min, diluted 2-fold in 1x G3 buffer (New England Biolabs) and incubated with 160 U endoglycosidase H (New England Biolabs) for 2h at 37°C. Concentrated NuPAGE sample buffer (Life Technologies) was added, and the digests were resolved by SDS-PAGE as described below. Ten percent of the remaining lysate was reserved for input for surface levels and the rest was diluted 5-fold in lysis buffer and mixed with 50 ul of streptavidin-agarose (high-capacity, ThermoFisher). After rotation at 4°C for 2h, biotinylated proteins were recovered from the beads after washing in lysis buffer by elution in gel sample buffer containing 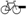 mercaptoethanol. Samples, including the endo H and surface input samples, were resolved on 4-12% NuPAGE gels (Life Technologies) in MES buffer. Following electrophoresis, proteins were transferred to PVDF for 90 min at 20 mA, and membranes were blocked with 5% nonfat milk in Tris-buffered saline (TBS) for 1h. Primary antibodies were incubated in Tris-buffered saline containing 5% nonfat milk and 0.05% Tween-20 overnight at 4°C. After washing, secondary antibodies (IRDye 800 donkey anti-rabbit IgG, LiCOR) were applied for 1h at RT and after washing, blots were quantified on LiCOR Odyssey CLx system and Image Studio (LiCOR). Analysis was in Prism GraphPad 8.0. The percent of protein at the cell surface was calculated in relation to that in the input sample (10% of that used for the surface sample).

### Flow cytometry for luminal Golgi pH determination

This was performed exactly as described [11], using the medial Golgi pHlorin, GnT1-pHlorin. Transfected cells were incubated in 100 ug/ml cycloheximide (Sigma) for 60 min prior to analysis to chase newly synthesized proteins from the endoplasmic reticulum. A standard curve was prepared from cells expressing the pHlorin alone, incubated in the presence of 10 uM monensin (Sigma) and 10 uM nigericin (Sigma) at pH 5.5, 6.0, 6.5, 7.0 or 7.5 before flow cytometry. Samples were excited at 405 and 488 nm and emission was collected with filters between 500 and 550 nm and 515 to 545 nm respectively, in a LSRII flow cytometer (Becton Dickenson). Then the fluorescence of cells expressing IBV M, IBV E or SARS CoV-1 3a along with the GnT1-pHlorin were measured in the absence of ionophores. Data were collected and analyzed using FACS Diva 8.0 software and Excel. The emission ratios of GnT1-pHlorin in buffers of different pH in the presence of ionophores was used to generate a standard curve for calculating the Golgi luminal pH of cells expressing IBV M, IBV E and SARS CoV-1 3a. Although the pandemic prevented repeat of this single experiment, over 3500 cells were analyzed and earlier work demonstrated that the results are quite reproducible [11].

## Results

### Golgi mophology is disrupted by SARS CoV-1 3a protein

To examine the effect of SARS CoV-1 3a overexpression on Golgi morphology, we transfected plasmids encoding the 3a protein (and IBV E and IBV M as positive and negative controls, respectively) into Vero cells. Of note, we did not observe any cell death (apoptotic or necrotic) in cells expressing 3a, as has been reported in other studies [24–26]. Perhaps initiation of cell death depends on higher levels of expression or on cell type. After 18 h, cells were fixed and stained for Golgi markers and the viral protein. Figure 1a shows the distribution of these proteins in transfected cells. SARS CoV-1 3a is transported past the Golgi to the plasma membrane, whereas IBV M and E are both retained in the Golgi region. Dispersion of Golgi complex proteins in cells expressing GnT1-pHlorin and the indicated viral protein was observed after staining with two Golgi markers: mouse anti-GM130 and rabbit anti-giantin (Figure 1b). Transfected cells were identified by the GFP signal from the pHlorin and indicated by white arrows. The Golgi was dispersed in cells expressing SARS CoV-1 3a, similar to cells expressing IBV E. Cells expressing the IBV M protein served as a negative control, since this protein is also targeted to the Golgi complex but does not induce dispersion.

**Figure 1.**
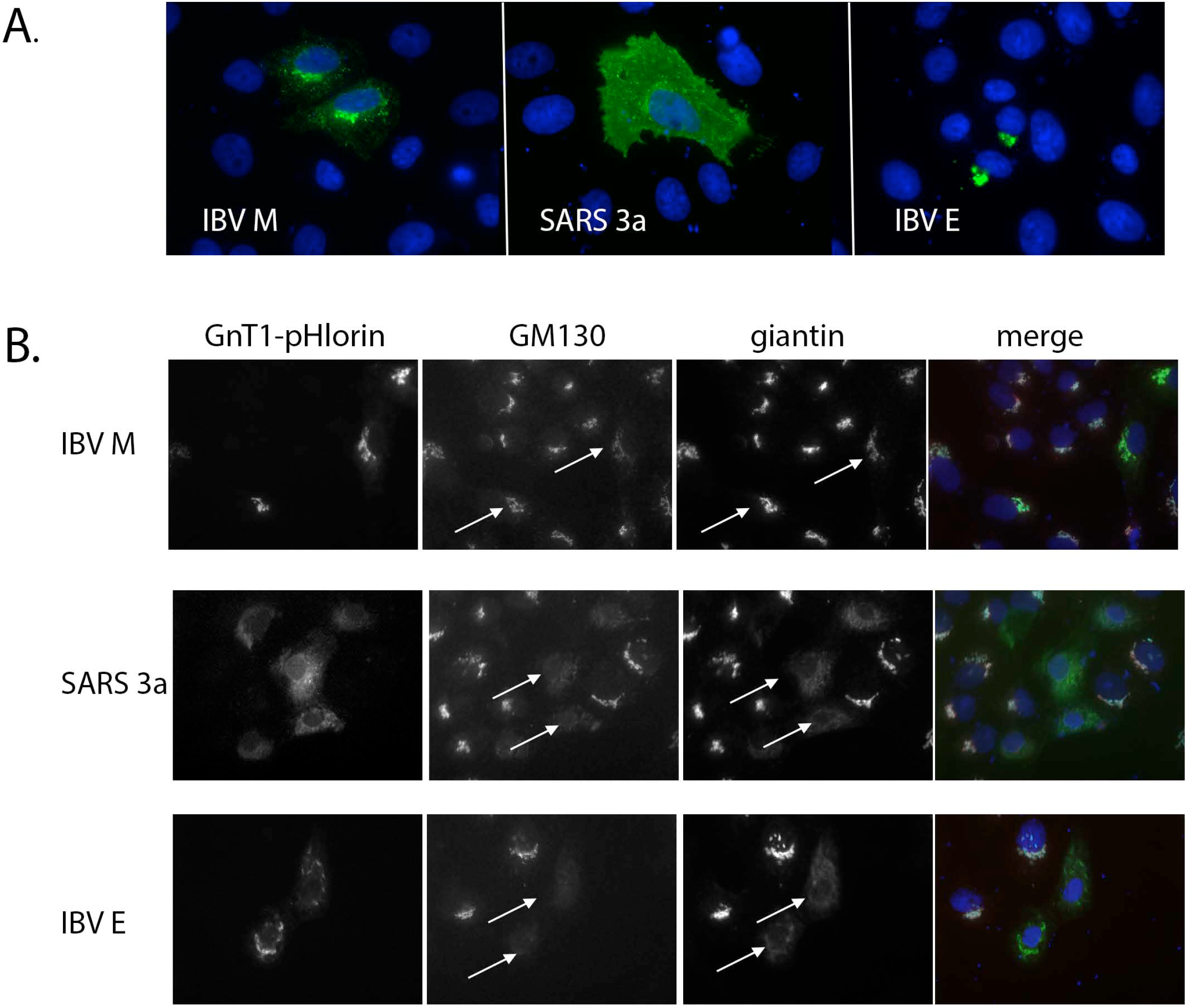
Golgi dispersion in cells expressing SARS CoV-1 3a protein. (**a**) Vero cells expressing IBV M, SARS CoV-1 3a or IBV E were stained with the relevant viral protein antibody (green). Nuclei are stained with Hoescht 33342 (blue); (**b**) Cells expressing IBV M, SARS CoV-1 3a or IBV E along with GnT1-pHlorin were stained with mouse anti-GM130 (red in merge), rabbit anti-giantin (cyan in merge) and Hoescht 33342 (blue in merge). Transfected cells (white arrows) were identified by GnT1-pHlorin expression. Golgi morphology is disrupted in cells expressing SARS CoV-1 3a and IBV E, but not IBV M.

To quantify these results, the area occupied by stained Golgi elements was determined using ImageJ in cells transfected and imaged as in Figure 1b. GM130 is a peripheral cis-Golgi protein and giantin is an integral membrane protein of the medial-Golgi. Golgi dispersion in cells expressing SARS CoV-1 3a protein was not as extensive compared to that in cells expressing IBV E (Figure 2). This might be because SARS CoV-1 3a moves through the Golgi to post-Golgi compartments, while IBV E is tightly retained in the Golgi region, as seen in Figure 1a. To test this idea, we attempted to generate a 3a mutant that was retained in the Golgi by swapping out the cytoplamic tail with that of SARS CoV-1 E, which contains Golgi targeting information [19]. However, this mutant was retained in the endoplasmic reticulum (possibly misfolded) so was not useful in addressing this possibility. The peripheral Golgi protein GM130 showed greater dispersion than giantin, possibly reflecting the release of peripheral proteins before Golgi membrane elements are dispersed.

**Figure 2.**
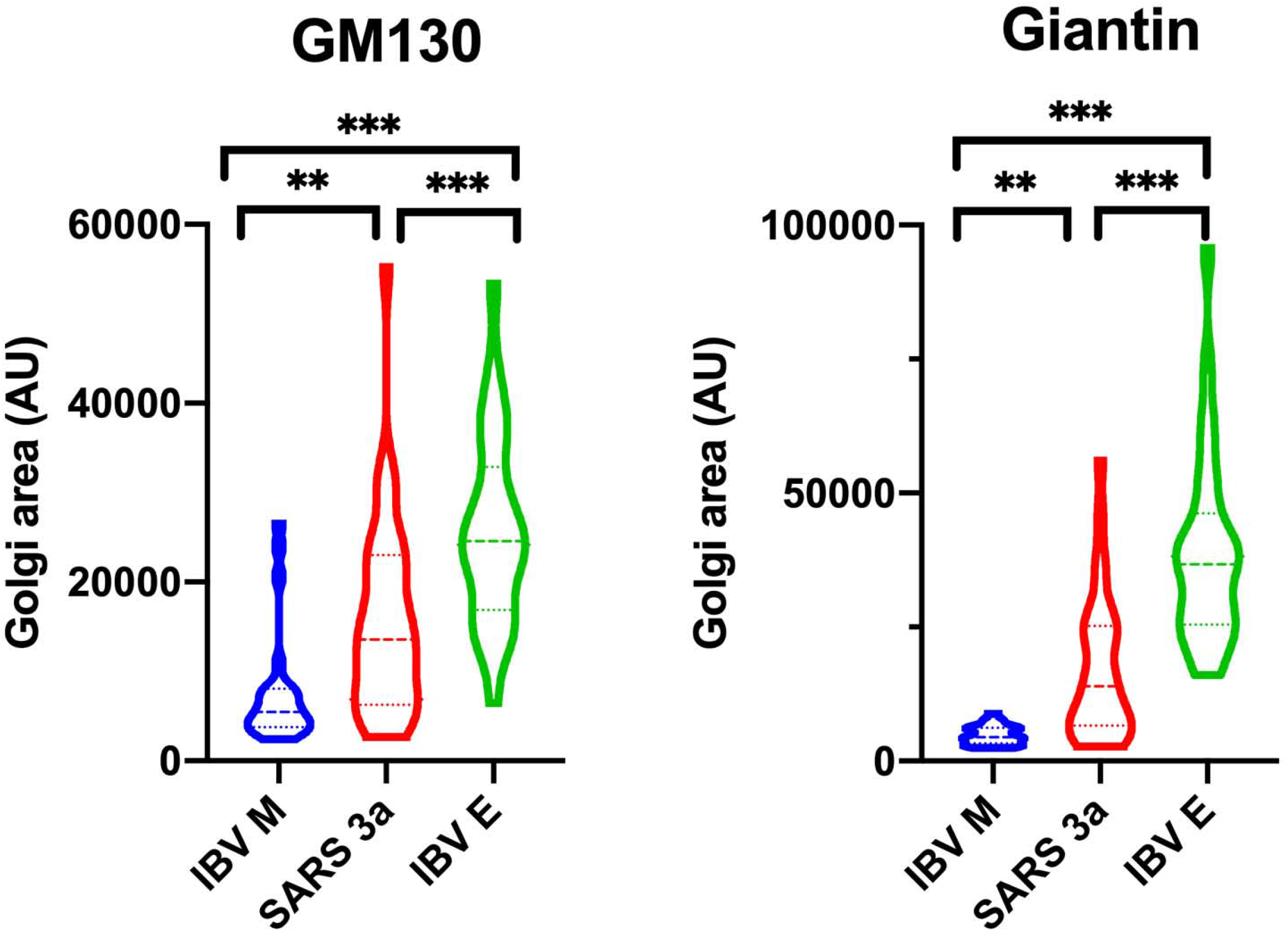
Quantification of Golgi dispersion. The area occupied by stained Golgi elements for both GM130 and giantin from two separate experiments in cells expressing IBV M, SARS CoV-1 3a or IBV E was quantified using ImageJ. Violin plots show median (dashed line) and quartiles (dotted lines). N= 24 (IBV M), 35 (SARS CoV-1 3a) and 26 (IBV E). Significance by one-way ANOVA and post-hoc Tukey test: *** p <0.001, ** p = 0.001-0.01.

### Cargo trafficking is reduced in cells expressing SARS CoV-1 3a

To determine if the morphological changes in the Golgi complex were accompanied by a reduced level of cargo traffic, we analyzed carbohydrate processing and surface levels of the vesicular stomatitis virus glycoprotein (VSV-G) in cells co-expressing SARS CoV-1 3a, IBV E or IBV M. The VSV-G protein is a commonly studied membrane protein with two N-linked oligosaccharides that are processed in the Golgi as the protein moves to the plasma membrane. At 20 h post-transfection, cells were first surface biotinylated using membrane impermeant NHS-sulfo-biotin. A portion of the lysate was reserved for digestion with endoglycosidase H (endo H) to determine the fraction of molecules that had been processed in the medial Golgi, and the biotinylated proteins in the remaining lysate were isolated on streptavidin-agarose. After SDS-PAGE and transfer to PVDF, VSV-G was identified by blotting with anti-VSV antibody. Representative blots are shown in Figures 3a and b, with quantitation of 3 experiments in Figure 3c. Both oligosachharide processing and surface expression for VSV-G were reduced in cells co-expressing SARS CoV-1 3a compared to the IBV M control, although not as strongly as in cells co-expressing IBV E. Reduced endo H processing and surface levels were also observed for cells co-expressing SARS CoV-1 S protein instead of VSV-G, to 46% and 28% of the negative control, respectively. The intermediate level of reduction of cargo trafficking compared to cells co-expressing IBV E paralleled the level of Golgi dispersion observed by indirect immunofluorescence microscopy, possibly reflecting the fact that the 3a protein is not retained in the Golgi like the IBV E protein.

**Figure 3.**
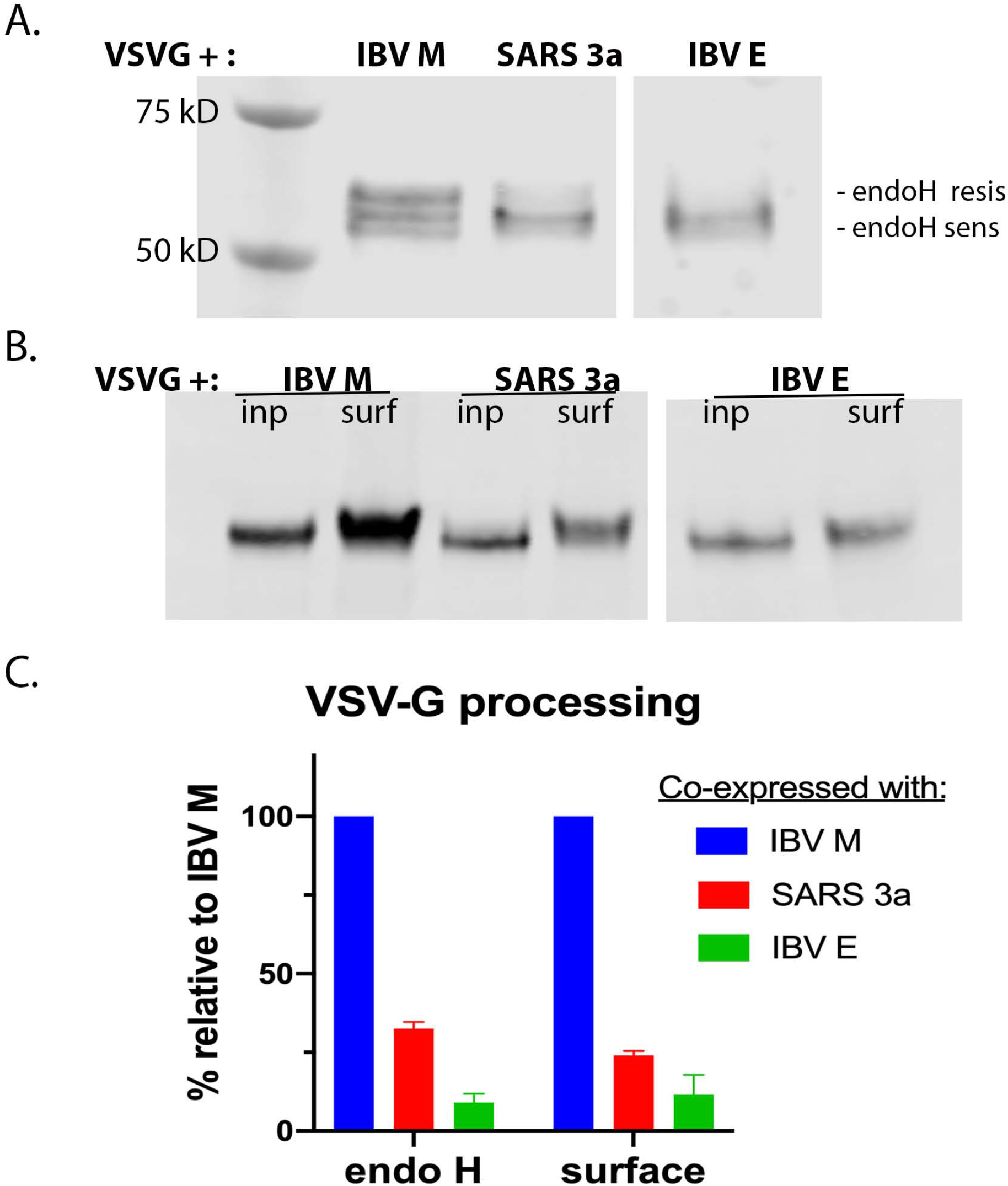
Cargo trafficking is reduced in cells expressing SARS CoV-1 3a. Vero cells co-expressing VSV-G and IBV M, SARS CoV-1 3a or IBV E were surface biotinylated. (a) A portion of the lysate was treated with endo H to determine oligosaccharide processing. (b) The remaining lysate was incubated with streptavidin-agarose to isolate surface proteins (after reserve of 10% for input (inp)). Panels (a) and (b) are representative blots and panel (c) shows quantification from 3 independent experiments. The negative control (IBV M) was set to 100% to allow comparison across multiple experiments. Values shown are mean ± SD.

### SARS CoV-1 3a protein neutralizes the Golgi complex

We used flow cytometry to analyze the fluorescence of a Golgi-targeted luminal pHlorin molecule (GnT1-pHlorin) as described earlier [11]. The fluorescence of the GFP derivative called pHlorin is quenched at acidic pH. Vero cells expressing only the GnT1-pHlorin were used to generate a standard curve after incubation in buffers at fixed pH in the presence of ionophores. Then cells co-expressing GnT1-pHlorin and SARS CoV-1 3a, IBV E or IBV M were subjected to the same analysis in the absence of ionophores. The ratio of fluorescence emission spectra was measured as described in Materials and Methods Figure 4a), and the pH of the Golgi lumen was calculated based on the standard curve. Although not as great a change as with IBV E, the pH of the Golgi lumen in cells expressing SARS CoV-1 3a was also increased (Figure 4b). We concluded that the Golgi dispersion and reduction in cargo trafficking was likely a result of the increase in Golgi pH.

**Figure 4.**
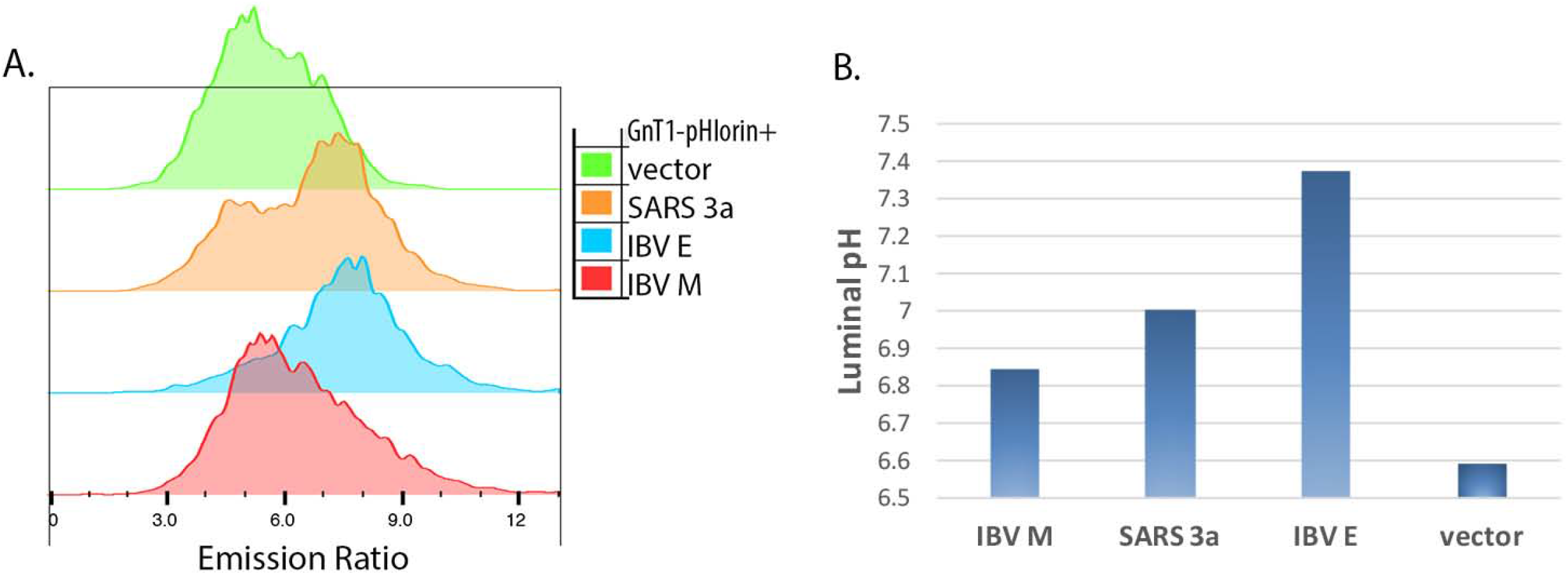
SARS CoV-1 3a neutralizes the Golgi. A standard curve was produced from cells expressing GnT1-pHlorin after incubation at fixed pH in the presence of ionophores. Flow cytometry was performed to determine the fluorescence emission ratios and >3500 cells were analyzed for each sample. (a) Emission ratios for cells expressing GnT1-pHlorin and either IBV M, SARS CoV-1 3a, IBV E or vector. (b) The standard curve was used to estimate Golgi luminal pH. Neutralization by the 3a protein was intermediate between IBV M and IBV E.

### The viroporin activity of SARS CoV-1 3a is important for Golgi disruption

We previously showed that the Golgi dispersion and neutralization by IBV E did not appear to require its viroporin activity since these phenotypes correlated with a monomeric form of E that cannot form an ion channel [11]. We speculated that interaction with a host protein that required a specific sequence in the IBV E transmembrane domain was the mechanism for the observed phenotypes. To test if the SARS CoV-1 3a protein required viroporin activity for Golgi dispersion, we generated several 3a proteins with mutations that were reported to inactivate ion channel function [16]. First, Y91 and H93 in the second transmembrane were replaced with alanines. This mutant protein was poorly expressed however, so we did not pursue its analysis. Next, a point mutation in the third transmembrane domain, Y109A, was generated. This protein was expressed as well as the wild-type protein and was distributed in a similar cellular pattern (Figure 5a). Golgi dispersion was less evident in cells expressing 3aY109A compared to wild-type protein. Quantification showed that the mutant protein did not disrupt Golgi morphology to the same extent as the wild-type 3a protein (Figure 5b). The same was found for cargo trafficking: for VSV-G endo H resistance, 3aYA was 66% of the negative control, and for surface, 88% (compare to Figure 3). Thus, unlike the IBV E protein, the SARS CoV-1 3a protein may disrupt secretory compartments directly through ion channel activity.

**Figure 5.**
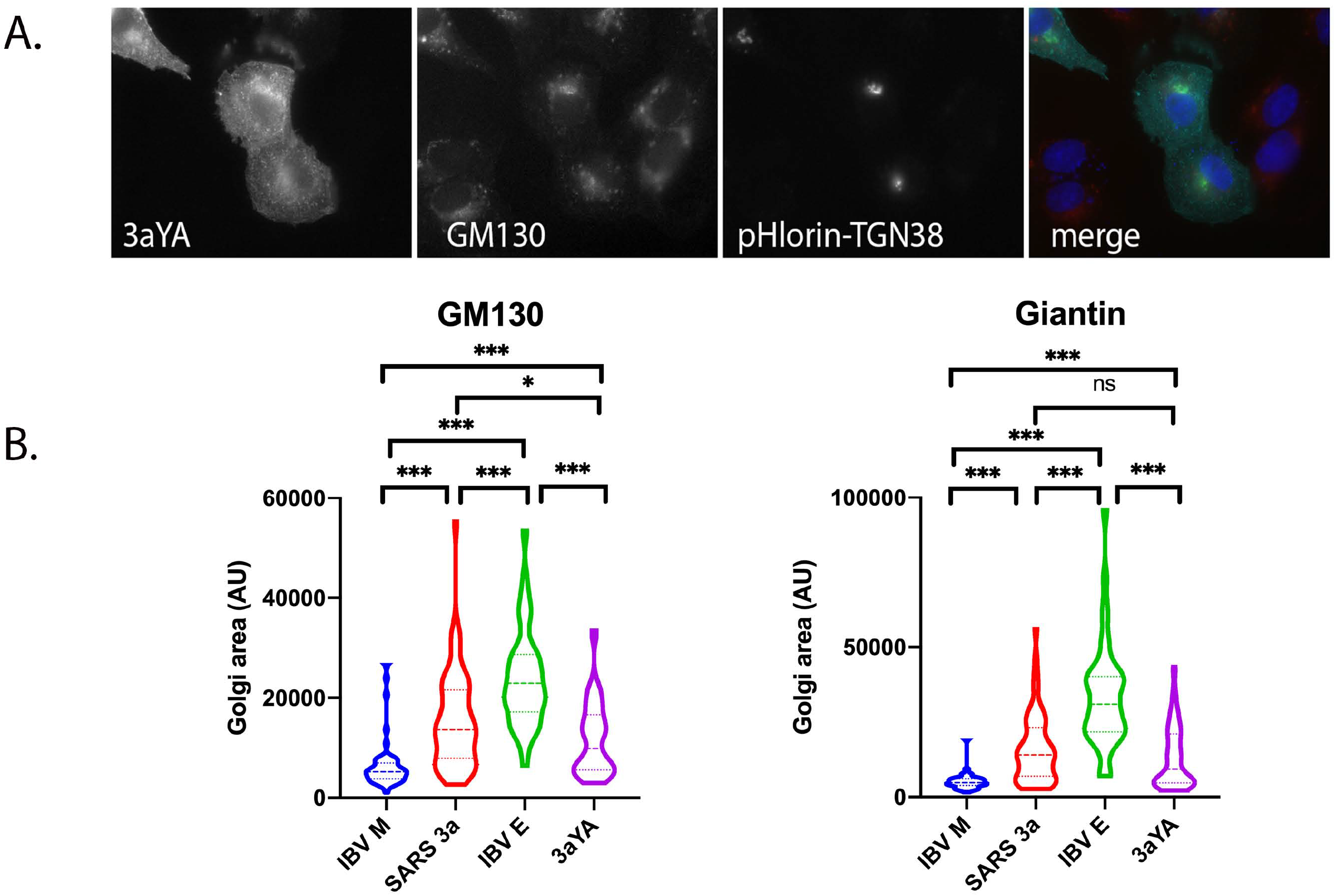
A mutation in 3a that abrogates viroporin activity *in vitro* reduces Golgi dispersion. (a) SARS CoV-1 3aY109A is distributed in a similar pattern as the wild type protein, but dispersion of Golgi elements is not as pronounced. (b) Quantification of Golgi area in 4 independent experiments as described in Figure 2. Violin plots show median (dashed line) and quartiles (dotted lines). N=56 (IBV M), 60 (SARS CoV-1 3a), 79 (3aY109A) and 39 (IBVE). Significance by one-way ANOVA and post-hoc Tukey test: *** p <0.001, * p = 0.01-0.05, ns, non-significant.

### SARS CoV-2 3a protein also fragments the Golgi

After the emergence of SARS CoV-2 that caused the COVID-19 pandemic, we were curious if the SARS CoV-2 E or 3a proteins might have similar effects on the Golgi complex. In cells overexpressing SARS CoV-2 E, we saw no morphological effects on the Golgi. However, overexpression of the SARS CoV-2 3a protein did induce alterations in Golgi morphology similar to those in cells expressing SARS CoV-1 3a (Figure 6). Golgi marker dispersion was observed for both the *cis*-Golgi GM130 protein and the *trans*-Golgi p230 protein. Even though the amino acid identity of SARS CoV-1 and -2 3a is only 73% (85% similar), the residues that are important for ion channel activity are conserved. Thus, it is likely that the mechanism for Golgi disruption for SARS CoV-2 is similar to that for SARS CoV-1.

**Figure 6.**
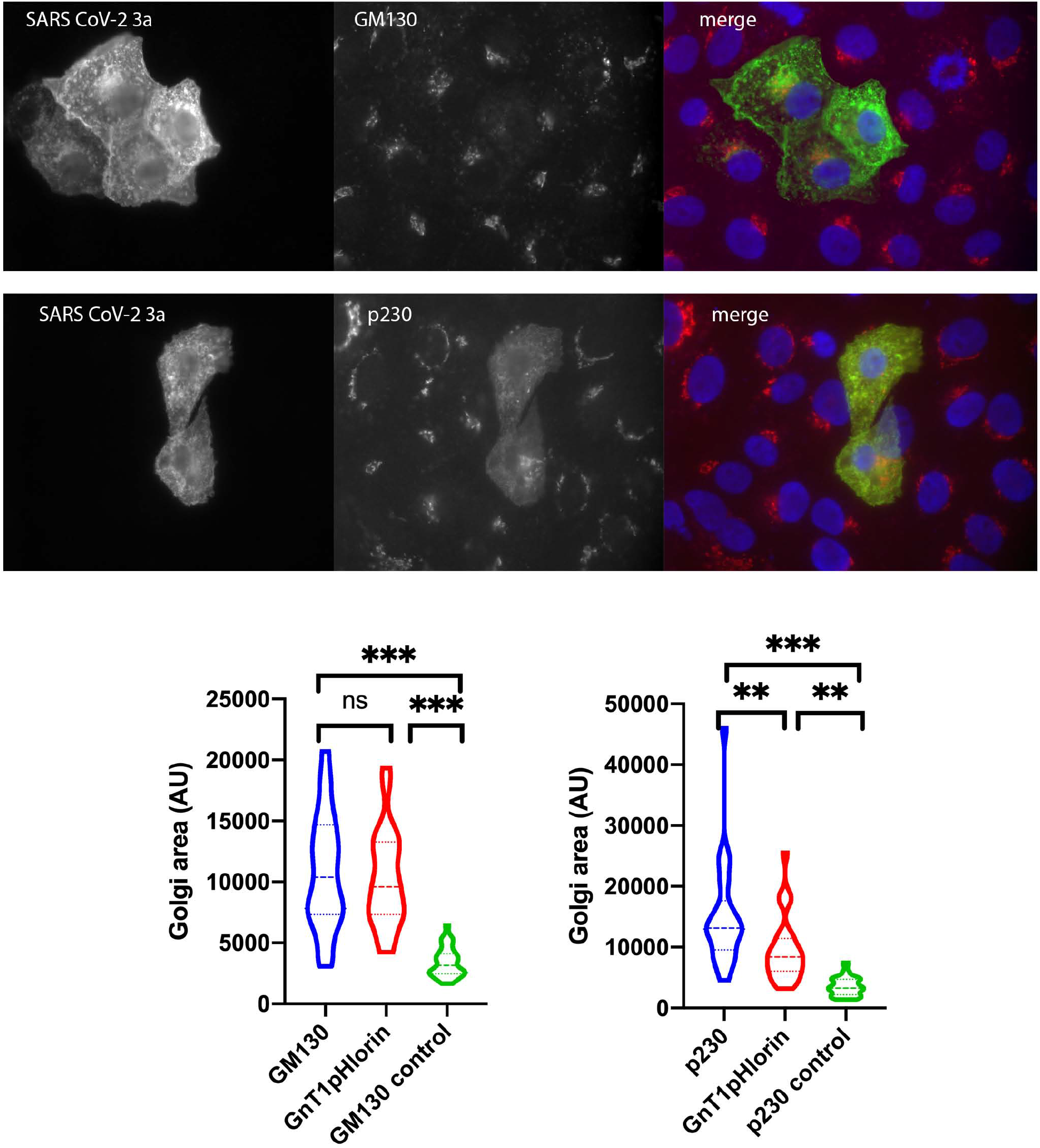
SARS CoV-2 3a protein alters Golgi morphology. Vero cells expressing SARS CoV-2 3a and GnT1-pHlorin were stained with mouse anti-GM130 or mouse anti-p230 (both red in merge) and rabbit anti-SARS 3a (green in merge) along with Hoescht 33342 (blue in merge). Both Golgi markers are dispersed to similar extents as in cells expressing SARS CoV-1 3a. The quantification of Golgi area in the lower panels was from two independent experiments, and the Golgi area in 3a-expressing cells was compared to that in neighboring non-transfected cells. Violin plots show median (dashed line) and quartiles (dotted lines). N= 23 (GM130), 26 (p230), 32 (non-transfected). Note the different scales when comparing GnT1-pHlorin. Significance by one-way ANOVA and post-hoc Tukey test: *** p <0.001, ** p <0.05, ns, non-significant.

## Discussion

The SARS CoV-1 3a protein has been implicated in virus pathogenicity [16], possibly through inflammasome activation and/or induction of cell death [24–27]. Ion channel activity is important for pathogenicity [16]. The SARS CoV-2 3a protein was recently reported to block autophagosome fusion with lysosomes and impair lysosome function, but interestingly SARS CoV-1 3a protein did not have this activity [28]. Whether this difference in SARS CoV-1 and -2 3a protein contributes to differences in infectivity is not yet known. Not all coronaviruses encode a 3a-like protein; a recent preprint reports that only pathogenic human coronaviruses that have their closest ancestors in bats have the protein, including SARS CoV-1, Middle East respiratory syndrome (MERS) virus and SARS CoV-2 [29].

We show here that the SARS CoV-1 3a protein disrupts Golgi morphology and cargo trafficking through the Golgi complex. We previously studied this phenotype induced by IBV, a gamma-coronavirus, and showed it was due to the E protein. IBV does not encode a 3a-like viroporin. However, overexpression of the SARS CoV-1 E protein did not disrupt the Golgi [9], leading us to investigate the 3a protein. The Golgi disruption in cells expressing SARS CoV-1 3a was not as dramatic as that in cells expressing the IBV E protein. We believe that this is due to the lack of retention of the 3a protein in the Golgi like the IBV E protein. The dispersion of Golgi markers, reduction in cargo trafficking and modest neutralization of the Golgi lumen is likely due to the transient presence of the 3a protein in this compartment. We attempted to construct a 3a protein that was retained in the Golgi to test this idea, but the chimeric protein may have been misfolded because it was not transported out of the endoplasmic reticulum and thus could not be used to address this possibility. We also showed that expression of SARS CoV-2 3a disrupts Golgi morphology, suggesting it may have similar properties to SARS CoV-1 3a.

A monomeric form of IBV E is responsible for the Golgi phenotypes and thus cannot be due to its viroporin function [11]. To test if SARS CoV-1 3a viroporin activity was required for Golgi disruption, we tested a mutant (Y109A) that was previously shown to eliminate ion channel activity *in vitro* [16]. Golgi dispersion in cells expressing this mutant protein was reduced relative to the wild-type protein, but not abrogated. It is possible that this single amino acid change is not enough to eliminate channel activity in cells. A recent cryo electron microscopy structure of SARS CoV-2 3a protein suggests that the main channel for ion transport is formed by transmembrane domains 1 and 2, whereas Y109 is in transmembrane domain 3 and may contact the neighboring dimer [29]. This may also be the case for SARS CoV-1 3a, since the proteins are relatively conserved. If so, this could explain why the Y109A mutant was not completely inactive. We also generated a 3a mutant with two substitutions in the second transmembrane domain (Y91 and H93 changed to alanines), which also lacks ion channel activity *in vitro* [16]. However, the protein was poorly expressed so was not used in our analyses.

Collapse of pH gradients in cellular organelles is known to disrupt membrane traffic and alter the composition and morphology of organelles [30, 31]. The disruption of Golgi function observed in cells expressing SARS CoV-1 3a (and IBV E) are likely due to neutralization of the Golgi complex. The mechanism of neutralization of acidic compartments by SARS CoV-1 3a could involve direct viroporin-induced collapse of potassium gradients across organellar membranes. Calcium may also be transported by the 3a channel [29], which could affect luminal proton concentrations directly or indirectly. For IBV E, we proposed that the monomeric form interacts with a host protein required for acidification (directly or indirectly) and inhibits its function [11]. Since IBV E is tightly retained in the early Golgi, this mechanism may make sense if downstream compartments need to be neutralized. By contrast, the SARS CoV-1 3a protein is trafficked to the plasma membrane and endo-lysosomal compartments so it could act directly to neutralize acidic compartments throughout the endomembrane system. It is interesting that a similar phenotype is present for two different coronaviruses but depends on different viral proteins and apparently different mechanisms. This suggests that neutralization of acidic compartments is an important feature of coronavirus-infected cells.

For a virus that assembles by budding into an early Golgi compartment and then must egress from the cell, reduction in cargo trafficking is unexpected. For IBV, neutralization protects the S protein from aberrant proteolysis as virions egress, keeping virus infectious [11]. Thus, the reduction in the rate of virus release may be an acceptable compromise. Neutralization of endosomes or lysosomes could also impact antigen presentation and contribute to immune evasion, as recently proposed [13].

One limitation of our study is that we only examined cells expressing the viral proteins from cDNA. In the future, it will be important to measure the luminal pH of Golgi, endosomes and lysosomes in infected cells. Our previous results with IBV E suggest that the phenotypes in cells overexpressing the protein might be exaggerated but accurately reflect the function in infected cells [7, 11, 12]. Proteolytic processing of S in both SARS CoV-1 and -2 infected cells when the 3a protein is compromised will also need to be assessed. Importantly, specific inhibition of 3a channel function with repurposed or new drugs may provide additional treatments for SARS, MERS and COVID-19 [3, 32].

## Author Contributions

Conceptualization, C.E.M.; methodology, R.R.G. and C.E.M..; formal analysis, R.R.G. and C.E.M.; investigation, R.R.G. and C.E.M. writing—original draft preparation, C.E.M.; writing—review and editing, R.R.G.; funding acquisition, C.E.M. All authors have read and agreed to the published version of the manuscript.

## Funding

This research was funded by the National Institutes of Health R01 GM117339.

## Acknowledgments

We thank Jason Westerbeck and Trisha Nilles for assistance with the flow cytometry, Stephen Gould and Chenxu Guo for the plasmids encoding SARS CoV-2 E and 3a and Yusuke Maeda for the pHlorin plasmids.

## Conflicts of Interest

The authors declare no conflict of interest.

## Notes

### Competing Interest Statement

The authors have declared no competing interest.

